# Transcranial Focused Ultrasound Remotely Modulates Extrastriate Visual Cortex with Subregion Specificity

**DOI:** 10.1101/2024.01.20.576476

**Authors:** Kai Yu, Samantha Schmitt, Yunruo Ni, Emily C. Crane, Matthew A. Smith, Bin He

## Abstract

Low-intensity transcranial focused ultrasound (tFUS) has emerged as a powerful neuromodulation tool characterized by its deep penetration and precise spatial targeting to influence neural activity. Our study directed low-intensity tFUS stimulation onto a region of prefrontal cortex (the frontal eye field, or FEF) of a rhesus macaque to examine its impact on a remote site, the extrastriate visual cortex (area V4). This pair of cortical regions form a top-down modulatory circuit that has been studied extensively with electrical microstimulation. To measure the impact of tFUS stimulation, we recorded local field potentials (LFPs) and multi-unit spiking activities from a multi-electrode array implanted in the visual cortex. To deliver tFUS stimulation, we leveraged a customized 128-element random array ultrasound transducer with improved spatial targeting. We observed that tFUS stimulation in FEF produced modulation of V4 neuronal activity, either through enhancement or suppression, dependent on the pulse repetition frequency of the tFUS stimulation. Electronically steering the transcranial ultrasound focus through the targeted FEF cortical region produced changes in the level of modulation, indicating that the tFUS stimulation was spatially targeted within FEF. Modulation of V4 activity was confined to specific frequency bands, and this modulation was dependent on the presence or absence of a visual stimulus during tFUS stimulation. A control study targeting the insula produced no effect, emphasizing the region-specific nature of tFUS neuromodulation. Our findings shed light on the capacity of tFUS to modulate specific neural pathways and provide a comprehensive understanding of its potential applications for neuromodulation within brain networks.

## Introduction

Transcranial focused ultrasound (tFUS) is a potent neuromodulation modality with high spatial specificity and deep penetration into the brain. It offers an opportunity for noninvasively modulating neuronal activity with high spatiotemporal resolution^1^, impacting neural connectivity, and even changing behavior outcomes. Rodent models have been widely used with a variety of neural recording approaches to explore the vast tFUS parameter space in pursuit of specific neuromodulation effects. However, using rodent models poses inevitable technical challenges for translational research and application due to the spatial and anatomical differences between rodents and humans. Through recent efforts, tFUS neuromodulation studies in nonhuman primates (NHP) have demonstrated that transcranial ultrasound can causally alter behavior (e.g., antisaccade (AS) latencies^2^ and choice behavior^3^) through cortical stimulation in FEF. Furthermore, tFUS stimulation can impact credit assignment and value representation by stimulating area 47/12o^4^ and anterior cingulate cortex (ACC)^4,5^, modulate regionally specific brain activity^6,7^, alter brain network coupling^4,6,8^, and further inhibit epileptiform discharge and reduce acute epileptic seizures^9^. Together, these studies point to the efficacy of tFUS as a neuromodulatory tool in large mammals.

Most studies examining the impact of tFUS on the brains of NHPs have measured blood-oxygen-level-dependent (BOLD) signal change using functional magnetic resonance imaging (fMRI) to assess whole-brain responses^4–8^. Although this recording modality provides a broad spatial measurement of the neurovascular signal across the brain, it does not provide direct measures of neural activity and faces challenges in capturing the fast brain dynamics^6^ in response to pulsed tFUS events due to its limited temporal resolution. Furthermore, measuring neurovascular signals in NHPs by fMRI limits the potential behavioral measurements that can be made, limiting potential observations of the outcomes of tFUS stimulation. These limitations on existing measurements of the impact of tFUS leave a substantial need for further work to better understand its impact as a tool for neuromodulation.

Our study aimed to measure the impact of tFUS in the NHP brain with high spatial and temporal resolution, incorporating extracellular recordings of neurons to capture neuronal activity as we manipulated tFUS parameters to examine its neuromodulatory effects. To do this, we leveraged a well-defined neural pathway from prefrontal cortex (FEF) to an extrastriate visual cortical area (V4)^10,11^, and measured neural activity (in V4) remotely from the ultrasound stimulation (in FEF). Our ultrasound neuromodulation leveraged a customized 128-element random array ultrasound transducer and employed the array’s focal steering capability to scan through stimulation targets.

This experimental design enabled us to demonstrate an impact of tFUS stimulation in FEF on V4 neuronal activity that was specific to the region of stimulation, consistent with the anatomical projections between FEF and V4. Overall, our work demonstrates the power of tFUS for noninvasively modulating neuronal circuits with subregional specificity and a clear impact on neuronal activity at remote sites.

## Material and Methods

### Experimental Model and Subject Details

We recorded from an implanted electrode array in the visual cortex of a rhesus macaque (*Macaca mulatta*) with concurrent tFUS stimulation. The subject was trained to perform a passive fixation task. The animal fixated centrally while a natural image was displayed for 400 ms in the V4 receptive field. After fixation, the animal performed a visually guided saccade to complete the trial (Fig. 2a). For each recording session, 120 trials with only visual stimulation (duration: 400 ms) and 120 trials with concurrent visual and tFUS stimulation were randomized and presented.

### Ultrasound Setup and Stimulation

We used a customized 128-element random array ultrasound transducer at a 700 kHz fundamental frequency transmitted from a relatively large acoustic aperture (diameter: 60 mm, F-number: 0.58). This ultrasound probe, called H275, was manufactured by Sonic Concepts, Inc. (Bothell, Washington, USA). A transducer holder (with X and Y axes indicators, using Tough 1500 Resin) and a customized ultrasound collimator (using Clear Resin) were 3D printed (Form 3+, Formlabs, Somerville, MA, USA) to fix the transducer’s location on the animal’s chair and assist the ultrasound targeting and steering along specific directions.

tFUS simulations based on a full-wave acoustic solver (Sim4Life, Zurich Med Tech ag., Zurich, Switzerland) were implemented in the presence of the NHP subject’s full skull model (based on CT images, see Supplementary Note 1: Structural Imaging). The customized ultrasound collimator was included in the computer simulations. With the subject’s MR-based brain model co-registered with the skull model, the tFUS beam and focus were moved onto the FEF region by adjusting the transducer’s position over the scalp. This relative positioning was further used as spatial guidance in the *in vivo* experiment to improve the precision of the tFUS targeting. In the sham ultrasound condition, the tip of the collimator was physically disconnected (with a 2-3 cm distance) from the prepared scalp surface while maintaining active ultrasound transmission. The X-axis of the H275 was aligned with the medial-lateral axis of the brain for focus steering.

### Intracranial Electrophysiological Recordings and Preprocessing

A 96-channel “Utah” electrode array (Blackrock Neurotech, Salt Lake City, UT, USA) with 1.0 mm length electrodes was implanted in visual cortical area V4 on the prelunate gyrus medial to the inferior occipital sulcus (see Stan et al. for more details of the array implantation in this animal RA^12^). The intracranial electrophysiological signals were acquired by a Ripple recording system and Trellis software suite at a sampling frequency of 30 kHz (Ripple Neuromed, Salt Lake City, UT, USA) with a bandpass filter of 0.3 Hz to 7.5 kHz set by the hardware for raw data acquisition, and additionally bandpassed with a range of 0.3 to 250 Hz for saving local field potentials. The local field potential data were further high-pass filtered during postprocessing in the FieldTrip toolbox with a cut-off frequency of 1 Hz and notch filtered at 60 and 120 Hz to remove powerline noise^13^. Spikes were obtained from this system by taking raw data, high-pass filtering (250 Hz to 7.5 kHz), and storing waveforms that exceeded a threshold set by a multiple of the root mean square noise on each channel. The resulting waveforms were manually sorted and examined in MKsort (Ripple Neuromed, Salt Lake City, UT, USA) after presorting with Spikesort toolbox (https://github.com/smithlabneuro/spikesort) to identify single units and multi-unit groups for further analysis (hereafter referred to as “neurons” for simplicity).

### Neural Electrophysiological Data Analysis

After the neural datasets were preprocessed, they were further segmented into epochs based on the visual/ultrasound stimulation events. For the LFP comparisons across different conditions, while all the epoch data were averaged across all the trials at specific representative channels in each set of experiments, individual trial data were retained to enable nonparametric statistical analyses using a temporal cluster-based permutation test with Monte-Carlo estimates of the significance probabilities in the FieldTrip toolbox^13,14^. The topographies of t-statistics at specific time frames with highlighted clusters of LFP change across all 96 channels were further generated in the toolbox by setting the significance level at 0.05. For the single-unit spiking analyses, peri-stimulus time histograms (PSTH) with a time bin size of 10 ms were used. The shaded areas behind the LFP mean profiles and the vertical lines on top of each time bin in the PSTHs represent +/− one standard error of the mean. Statistics for comparing the changes of single-unit spiking rates were produced with Wilcoxon rank sum tests after false discovery rate (FDR) corrections for multiple comparisons.

## Results

### Characterization of a random array-based transcranial focused ultrasound targeted at FEF

Single-element ultrasound transducers have been employed previously to modulate brain activity in NHPs^3–9^, but this technology has a major compromise in axial specificity. Such transducers need mechanical movement to adjust the spatial locations of the ultrasound focus if one would like to scan spatially across a targeted brain region. The customized spherically-concave, highly focused 128-element random array ultrasound transducer (H275, element diameter: 4 mm) working at a fundamental frequency of 700 Hz (-3 dB operating band: 648 – 777 kHz) was employed to achieve electrical spatial beam steering capability with improved focal performance. The diameter of its acoustic aperture is 60 mm with a concave radius of 35 mm. This customized focused ultrasound (FUS) probe (Fig. 1a, the random element spatial configuration as illustrated in the inset) was driven by a Vantage 256 research ultrasound platform to transmit FUS pulses through the skull. By simulating the focused ultrasound in free water, the measured coherent axial and lateral resolutions (-6 dB contour width and length) are 5.61 mm and 1.89 mm, respectively (Fig. 1a-c), and both dimensions were physically measured as 6.13 mm and 2.15 mm, respectively in degassed water (Fig. 1i-j). In the presence of a fully hydrated skull sample piece (from a different rhesus macaque than was used in this study) immersed in degassed water, the transcranial ultrasound focus was still spatially tight with a lateral focal size of 2.67 mm and an axial focal size of 7.71 mm (-6 dB contour width and length) (Fig. 1k-l). The acoustic insertion loss due to the skull was measured as approximately -9.1 dB.

**Figure 1.**
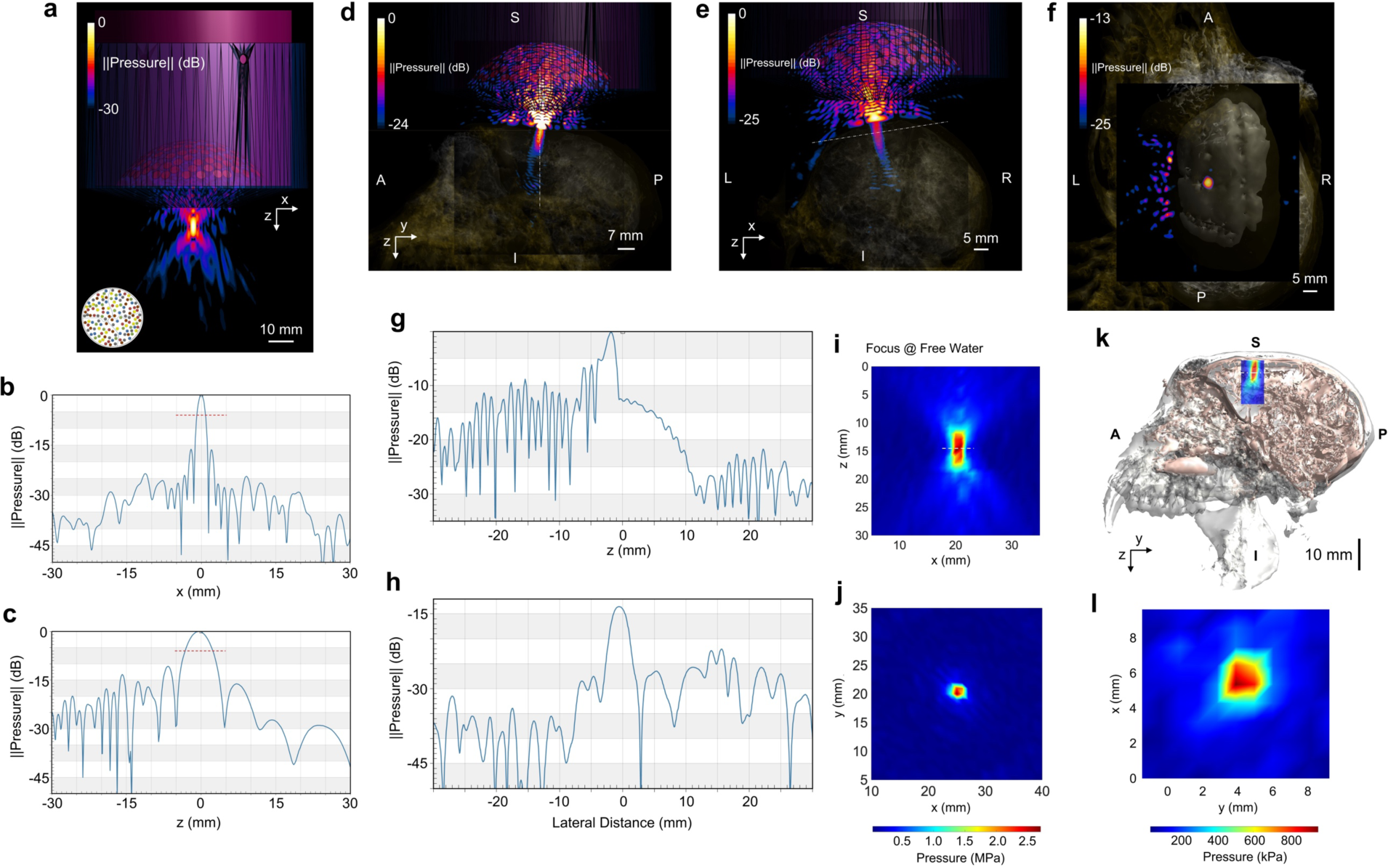
The characterizations of the ultrasound pressure field generated by the 128-element random array ultrasound transducer H275 in free water (**a-c**, and **i**-**j**) and in the presence of the nonhuman primate’s skull (**d**-**h**, and **k**-**l**). **a**, The 128-element ultrasound random array simulation with the presence of a customized 3D-printed collimator (purple solid) in free water. The random array distribution is illustrated in the inset. **b**-**c**, The line profiles of the ultrasound pressure field along the lateral (**b**) and the axial (**c**) directions. The dashed red lines indicate the -6 dB width. **d**-**f**, The transcranial simulation results in the sagittal (**d**) and coronal (**e**) in the presence of the nonhuman primate (NHP) subject-specific skull and brain models. **f**, A lateral characterization of the transcranial focus at the frontal eye field. **g**-**h**, The line profiles of the ultrasound field along the depicted dashed white lines in (**d**) and (**e**), respectively. **i**-**j**, The physical ultrasound focus of the real H275 device as measured with a hydrophone and reconstructed in free water. **k**-**l**, The physically measured transcranial ultrasound focus in the axial plane (**k**, co-registered with the NHP skull and brain models) and in the lateral plane (**l**) in the presence of a fully hydrated NHP skull sample.

We further simulated the pressure field in the presence of a complete subject-specific skull and brain models when the ultrasound stimulation was targeted at FEF. A sagittal view of the tFUS pressure field is depicted in Fig. 1d, with the central dashed white line indicating the location of the spatial profile as shown in Fig. 1g. The transcranial ultrasound axial specificity was 6.90 mm (-6 dB width), slightly better than the measured size in the physical phantom scanning. In the coronal view as shown in Fig. 1e, we took a line profile (dashed white line) perpendicular to the tFUS beam, and it showed a simulated lateral focal size of 3.18 mm (-6 dB width, Fig. 1h). This value was larger than the physical measurement of the lateral focal size of 2.67 mm in the presence of a skull sample. A 2D lateral view of the tFUS focus with the co-registered brain is further illustrated in Fig. 1f. The transcranial ultrasound focus was localized at the anterior bank of the arcuate sulcus, where FEF is typically found, based on the MR-based brain model. The dimensional characterizations of the transcranial ultrasound focus generated by the H275 enabled further tests on region-specific neuromodulation effects on FEF.

### tFUS remotely modulating the V4 activities in response to visual stimuli

FEF is a key player in the generation of eye movements and in coordinating the visual and oculomotor systems, and relatively low electrical current intracortical stimulations can reliably trigger saccadic eye movements^15^. Kubanek et al. demonstrated that tFUS stimulation at either left or right FEF changed saccadic directional biases and, thus, choice behaviors^3^. Our work builds on this study in two key ways: (1) we used more focused tFUS stimulation with electronic beam steering capability and (2) we sought to determine the impact of ultrasound stimulation on neural activity remote from the stimulation site. This latter difference is essential to the using tFUS as a modulatory stimulation to impact interaction among brain regions – in our case, by modulating activity in FEF to impact its interactions with visual cortex. To measure the impact of stimulation, we leveraged the known connections between FEF and V4 (for review, see Moore et al.^16^). During ultrasound stimulation of FEF, we recorded LFPs and multi-unit activities (MUA) from V4 (Fig. 2a). We did this because (1) it permitted large scale recordings to measure the impact of stimulation without a direct impact of stimulation on the recorded tissue, (2) it permitted a test of how well ultrasound stimulation could modulate a known circuit that is engaged in normal behavior and has been tested with electrical microstimulation^16^. We performed stimulation in the context of a simple fixation task with a visual stimulus (a natural image; the same image was used on every trial). The image was presented to increase the firing rate of the neuronal population and enable better detection of the modulatory impacts of ultrasound, which might take the form of suppression or enhancement. For each experimental session, we randomized trials in which ultrasound stimulation was performed simultaneously with the visual stimulus presentation (Fig. 2b) and trials in which the visual stimulus was presented alone without ultrasound stimulation.

**Figure 2.**
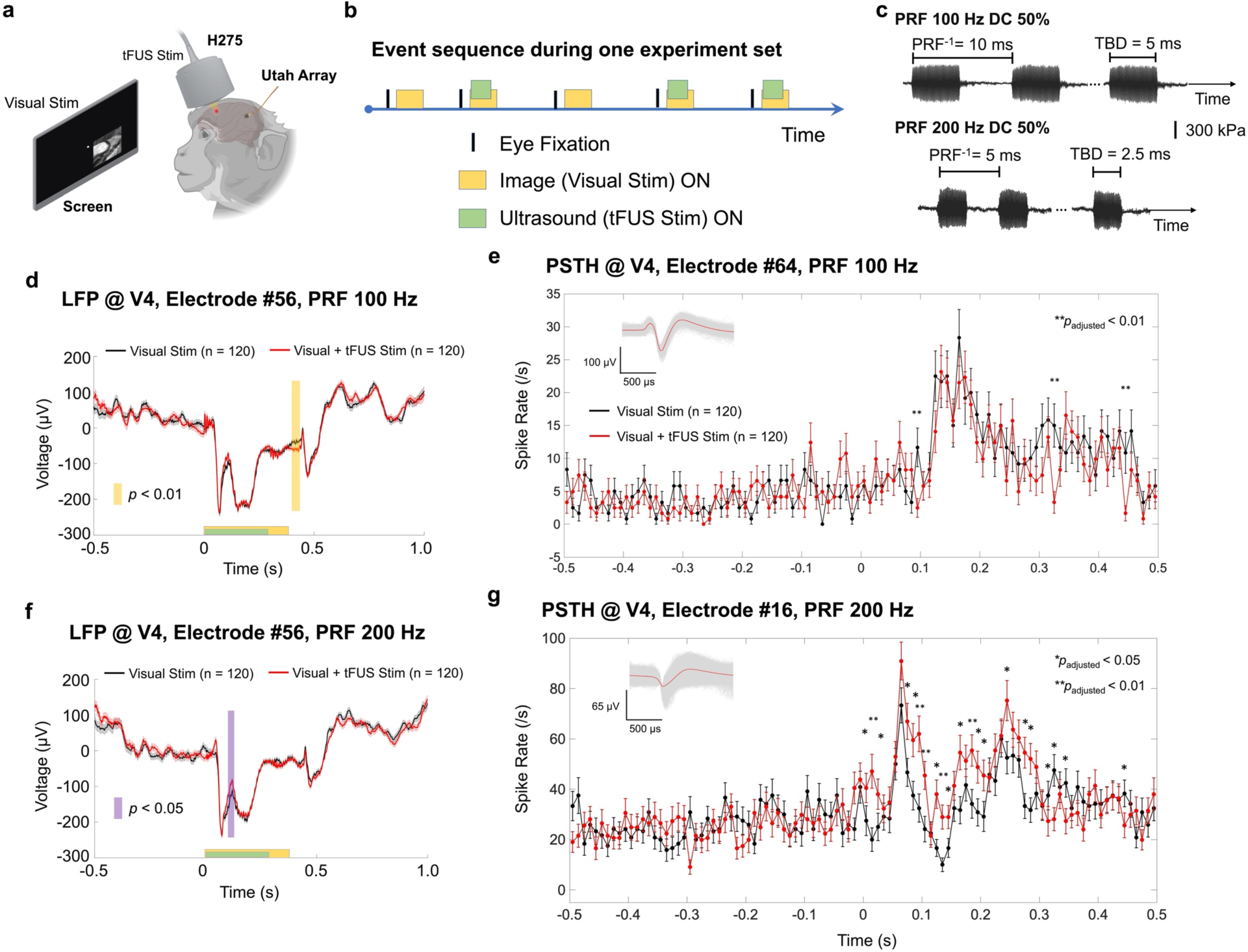
**a**, The experimental setup for intracranial neural recordings through a Utah array implanted in area V4 and recording during visual only or visual-tFUS stimulations. The tFUS stimulation was targeted at the frontal eye field (FEF). **b**, Event sequence during one experiment set with tFUS randomly occurring during half of the trials. The visual stimulation was delivered during fixation. **c**, The transcranial ultrasound temporal profiles applied in the *in vivo* experiment employed a pulse repetition frequency (PRF) of 100 Hz or 200 Hz with the same duty cycle (DC) of 50%. **d**, The local field potential (LFP) recorded from Electrode #56 at V4 area showed significant differences between the visual stimulation only (solid black line as the mean, and gray shaded area as the S.E.M.) and the hybrid stimulation (solid red line as the mean, and pink shaded area as the S.E.M.) with the tFUS stimuli of 100 Hz PRF and 50% DC. The statistics are obtained from a cluster-based nonparametric permutation test. The timing of visual/tFUS (n = 120) stimulations are depicted as the yellow and green boxes, respectively. **e**, The peri-stimulus time histograms (PSTHs) of an example single unit recorded from Electrode #64 (action potential waveforms sorted in the inset) in response to the visual only (black) and the hybrid stimuli (red). The time bins with significantly decreased spike rates are denoted with \*\**p_adjusted_* < 0.05 after the onset of the stimulus, i.e., 0 – 500 ms. Statistics with one-sided Wilcoxon rank sum test were corrected with a false discovery rate (FDR) for multiple comparisons. Error bars represent ± one S.E.M. at individual time points. **f**, LFP recorded from Electrode #56 in area V4 showed significant differences between the visual stimulation only (solid black line as the mean, and gray shaded area as the S.E.M.) and the hybrid stimulation (solid red line as the mean, and pink shaded area as the S.E.M.). **g**, The PSTHs of a neuron recorded from Electrode #16 (action potential waveforms sorted in the inset) in response to the visual only (black) and the hybrid stimuli (red). The time bins with significantly changed spike rates are denoted with \**p_adjusted_* < 0.05 or \*\**p_adjusted_* < 0.01 after the onset of the stimulus, i.e., 0 – 500 ms. Statistics with two-sided Wilcoxon rank sum test are corrected with FDR for multiple comparisons. Spike data at each time point are shown with mean±S.E.M.

To test the neuromodulation effects, we recorded the LFP and MUA from V4 when two types of tFUS stimulation (ultrasound waveforms and core parameters illustrated in Fig. 2c) were delivered to FEF. Both stimulation types shared the same duty cycle (i.e., 50%). The first stimulation type used a 100 Hz pulse repetition frequency (PRF) with a tone burst duration of 5 ms. The LFP recorded from one electrode in area V4 during the hybrid stimulation condition (i.e., visual stimulation with simultaneous tFUS stimulation) showed a statistically significant difference during the time window from 400 to 443 ms (Fig. 2d), starting 100 ms after the end of sonication. Specifically, comparing the peri-stimulus time histogram (PSTH) in visual stimulation with that in the hybrid stimulation (Fig. 2e), an example single neuronal unit significantly reduced its spiking rates at 15, 335 and 455 ms (\*\**p_adjusted_* < 0.01). Thus, this comparison demonstrated a potential inhibitory effect of the 100 Hz PRF tFUS at the FEF on the visual evoked response remotely at the V4. At the same LFP recording electrode as shown in Fig. 2d, 200 Hz PRF tFUS produced an excitatory effect during the sonication period, and significant differences in the LFP waveforms were seen during 92 – 131 ms (Fig. 2f). Consistent with this LFP finding, the PSTHs for another neuron (Fig. 2g) exhibited significantly increased spiking activities (\**p_adjusted_* < 0.05, \*\**p_adjusted_* < 0.01) at multiple time points during sonication, while exhibiting significantly reduced spiking rates after ultrasound stimulation ceased. The waveforms of the corresponding neuronal action potentials are shown in the inset of Figs. 2e and 2g, respectively. In summary, we observed that, when applied to FEF, different ultrasound PRFs with the same duty cycle induced bidirectional neuromodulation effects remotely at V4.

### The remote modulation on the V4 activities is subregional specific

With electrical stimulation, eye movement amplitude and direction can be controlled by the spatial location of stimulation within FEF^15^. That is, FEF contains a spatiotopic map of eye movement directions. Electrical stimulation of FEF also has a retinotopic-specific effect on V4 – the spiking response of individual neurons is modulated by stimulation of FEF only if that stimulation is matched to the V4 receptive field^16^. We used this property to determine the efficacy of our spatial steering of ultrasound stimulation at FEF. In Fig. 3a-b, the transcranial ultrasound focus of H275 from physical measurements is overlaid with the NHP brain model, and we targeted the ultrasound focus to the anterior bank of the arcuate sulcus based on focused ultrasound simulations incorporating the subject-specific MRI and CT data. Our goal was to verify the accuracy of spatially steered tFUS focus in our experimental paradigm. Our physical measurements showed that the steered tFUS focus was intact without significant aberration (Fig. 3c). The spatial locations of the steered tFUS focus were moved accurately along the X, Y and Z directions, and Fig. 3c shows the tFUS steering along the X and Z directions. To maintain the same ultrasound pressure level during steering, the focal pressures were compensated by adjusting the transmission power control of the Vantage system accordingly.

**Figure 3.**
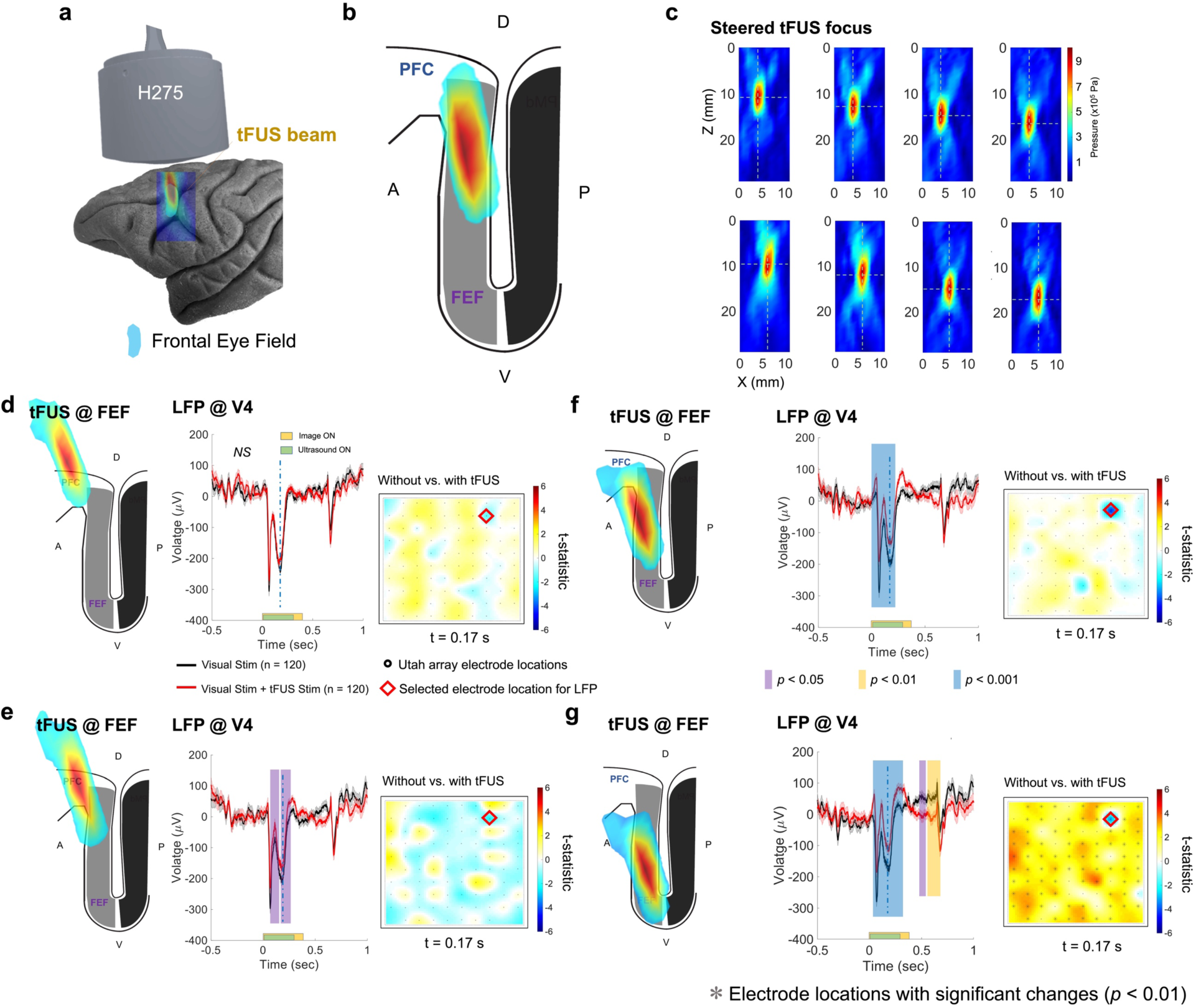
The tFUS remote neuromodulation effects on the visual-evoked potential (VEP) in the LFP are subregional specific. **a**-**b**, Registering the location and size of tFUS focus generated by H275 with the FEF region of the NHP brain model. **c**, The physically measured ultrasound focus behind a sample NHP skull during the electronic beam steering along the depth (Z, in rows) and the lateral direction (X, in columns). The dashed cross is located at the spatial peak point of the individual transcranial pressure field. **d**-**g**, When the ultrasound focus was steered along the depth of FEF, the tFUS (2 kHz PRF and 50% DC) neuromodulation effect on the VEP was more robust with increased significant changes during and after the sonication period. The statistics shown are based on nonparametric cluster-based permutation tests. All the plots depict the LFP acquisitions (n = 120 trials for each condition) from the same recording channel of the Utah array (denoted with a red diamond in the color map of each panel). The color maps indicate the statistical differences between spatial cluster topographies across 96-channel recording sites over the V4 at 0.17 s (indicated with vertical dashed lines in the LFP time voltage traces). The purple, yellow and blue vertical bars represent the time windows of significant changes with *p* < 0.05, < 0.01, and < 0.001, respectively. Statistics are calculated from permutation-based Monte-Carlo estimates of the significance probabilities with cluster-based correction for multiple comparisons.

Furthermore, we set out to test the subregional specificity of tFUS neuromodulation at FEF. We hypothesized that by adjusting the spatial location of the ultrasound focus across the anterior bank of the arcuate sulcus, the remote neuromodulation effects at V4 would change accordingly. Initially, at superficial locations denoted as Δz = 0 mm, adjusting the lateral locations of tFUS focus by ±2 mm (i.e., Δx = ±2 mm) did not modulate the visual-evoked LFP at the selected electrode in V4 (Fig. 3d, the t-statistic topographic map shows the differences across all 96 channels in V4 with a red diamond shape indicating the selected electrode location of the presented LFP). This indicates that the tFUS beam may not have reached FEF or modulated it effectively at this superficial location. However, once the focus was steered in the ventral direction (Δz = 2 mm, Δx = 2 mm), significant differences were observed in the LFP waveforms at the time windows of 60-134 ms and 156-263 ms (Fig. 3e) during the sonication period. We infer that the transcranial ultrasound focal spot started overlapping with FEF at this depth. By further directing the focus more ventrally (Δz = 4 mm) without implementing lateral steering, a neuromodulation effect with a higher level of statistical significance was detected during the time window of 30-317 ms, throughout most of the sonication period (i.e., 1-300 ms). More extensive neuromodulation effects were also seen beyond the sonication period when the ultrasound focus reached an even deeper location in the arcuate sulcus (Δz = 6 mm). Significant differences in the LFP waveforms with and without tFUS were observed during the time windows of 32-342, 472-547 and 555-698 ms. The right panels of Fig. 3d-g illustrate the t-statistical maps across the electrode array when comparing all LFP waveforms in visual stimulation with those in the hybrid stimulation at 110 ms after stimulation onset. The middle panels of Fig. 3d-g present the LFP waveforms acquired from the selected electrode location indicated by the red diamond shape. Overall, we saw that the remote neuromodulation effects depended on the spatial location of ultrasound focus within the arcuate sulcus.

### Remote modulation of V4 activity by ultrasound is parameter-dependent

In our experiment, the recordings were remote from the direct impact of ultrasound stimulation, helping avoid the possibility that any physical artifact could induce changes in recorded neural activity. To illustrate that the effect of ultrasound in our experiments occurred through modulation of neural activity in FEF and not some other nonspecific sources, we also performed a sham ultrasound study by physically disconnecting the acoustic aperture from the animal scalp while keeping the ultrasound transmission. In these experiments, the remote modulation effect was abolished entirely (Fig. 4a). When delivering 1 kHz PRF tFUS stimulation onto the FEF, a significant difference (*p* < 0.05) was observed during the sonication period in the LFP waveform (Fig. 4b). By increasing the ultrasound PRF to 2 kHz while maintaining the duty cycle at 50%, a much higher level of significant difference (*p* < 0.001) was noticed during sonication. The tFUS neuromodulation effect was also seen during a post-sonication time window (Fig. 4c). By further doubling the PRF to 4 kHz with the same duty cycle, extensive tFUS-induced changes in the LFP waveforms were observed during the sonication period (Fig. 4d). Thus, at this site, a PRF-specific effect of neuromodulation was observed that was absent during sham stimulation.

**Figure 4.**
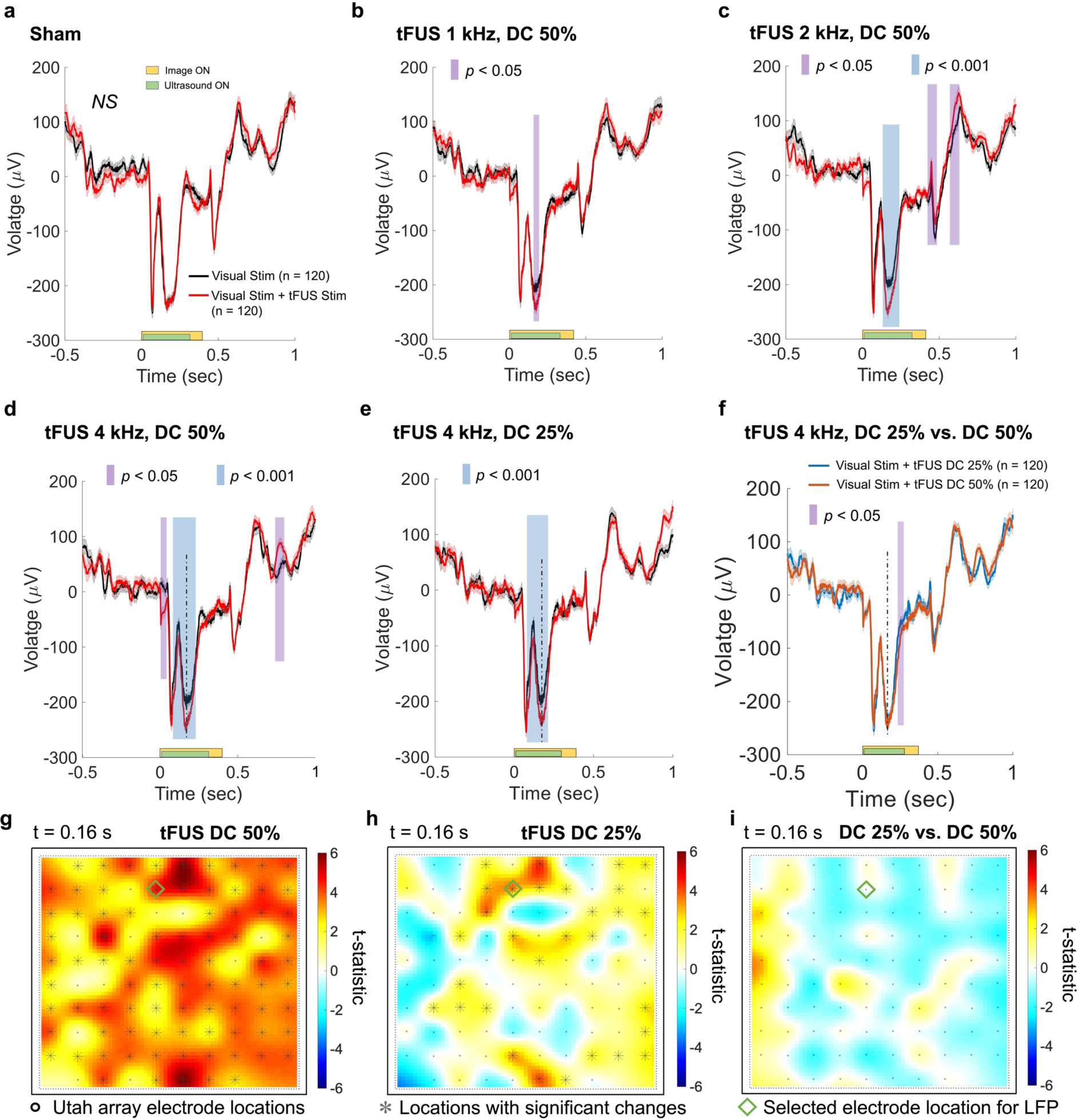
The tFUS remote neuromodulation effects on the VEP in the LFP are parameter dependent. **a**, In the sham ultrasound condition (i.e., decoupling the active sonication aperture from the NHP scalp), no significant difference was observed in the LFP comparison. **b**, A brief VEP change was observed at V4 when tFUS (1 kHz PRF and 50% DC) was administered to FEF. **c**, Doubling the ultrasound PRF (maintaining the same DC) while remaining at the spatial location of ultrasound focus at FEF enhanced VEP changes during sonication and further induced post-sonication changes. **d**-**f**, Comparisons on temporal features of LFP waveforms modulated by 50% DC (**d**) and 25% DC (**e**) while maintaining the same ultrasound PRF at 4 kHz. A slight difference (*p* < 0.05) was observed at the end of the sonication period, while the major component of the VEP in the middle of the sonication period (t = 0.16 s, indicated by the vertical dashed line) did not change significantly (**f**). **g**-**i**, Spatial cluster analyses of topographical differences across the 96-channel recording sites over the V4 remotely modulated by tFUS stimulation on FEF at the 50% DC (**g**) or the 25% DC (**h**) at 0.16 s after stimulus onset. Significantly different clusters (significance level: 0.05) are denoted with gray asterisks at corresponding electrode locations. At 0.16 s, no significantly different cluster was observed when directly comparing the topographies from DC 50% vs. DC 25% (**i**). Statistics are calculated from permutation-based Monte-Carlo estimates of the significance probabilities with cluster-based correction for multiple comparisons. The green diamond depicts the location of the selected electrode where the presented LFPs were recorded.

Duty cycle is another important sonication parameter that potentially changes neuromodulation effects^17–19^. When we decreased the ultrasound duty cycle to 25% while maintaining the same PRF level at 4 kHz, we continued to observe that differences between the conditions with and without tFUS remained (Fig. 4e). A direct comparison between the LFP waveforms obtained from those two duty cycle conditions (keeping the same PRF) showed some brief differences from 204 to 256 ms (Fig. 4f). The negative peaks at 0.16 s (indicated with dashed vertical lines) are not significantly different from each other (Fig. 4i) across all 96 channels of the V4 array, according to a spatial cluster analysis. The spatial cluster analysis (presented in Fig. 4g-h) demonstrated that at the same time point (e.g., 0.16 s), the higher duty cycle of tFUS led to a broader impact on V4 recordings across the area, possibly due to an increased deposition of ultrasound energy. The green diamond shape in the t-statistic topographic map indicates the selected electrode location where the LFP waveforms in Fig. 4a-f were recorded.

### tFUS remotely elicits V4 activity without visual stimuli

Until this point, the tFUS remote neuromodulation effects we assessed were in the presence of visual stimulation of the V4 neurons we recorded. This choice is aligned with previous work with electrical microstimulation of FEF, where the effects of stimulation were the strongest when V4 was driven by a visual stimulus^16^. Here, we set out to test whether the sole tFUS stimulation at FEF would elicit V4 activity without visual stimulation. As illustrated in Fig. 5a, a dot at the center of the screen was presented to the animal for eye fixation before the tFUS stimulation was delivered. The task for the animal (passive fixation followed by a saccade) and the duration of each trial were the same as above, except no visual stimulus was shown (other than the fixation dot). After repeating the ultrasound trials randomly in half of all 240 trials, the recorded LFP from one electrode at V4 showed significant differences during the time window from 144 to 192 ms (Fig. 5b). This demonstrates that without presenting an explicit visual stimulus, the applied tFUS (PRF: 3 kHz, Duty cycle: 60%) was able to evoke remote neural activity in V4 directly. In Fig. 5b, a red diamond indicates the location of the selected electrode for the LFP. The t-statistic topographic map at 0.16 s comparing the LFPs across the whole electrode array illustrates the differences between with and without tFUS stimulation (Fig. 5c). When compared to the time-frequency representations of the trials without ultrasound stimulation (Fig. 5d), the tFUS trials showed significantly increased spectral contents in the beta band (13-30 Hz, Fig. 5e) from 17.6 to 27.6 Hz during 11-170 ms, and from 24.3 to 29.6 Hz in the time window of 571-640 ms after sonication (Fig. 5f, -6 dB contour).

**Figure 5.**
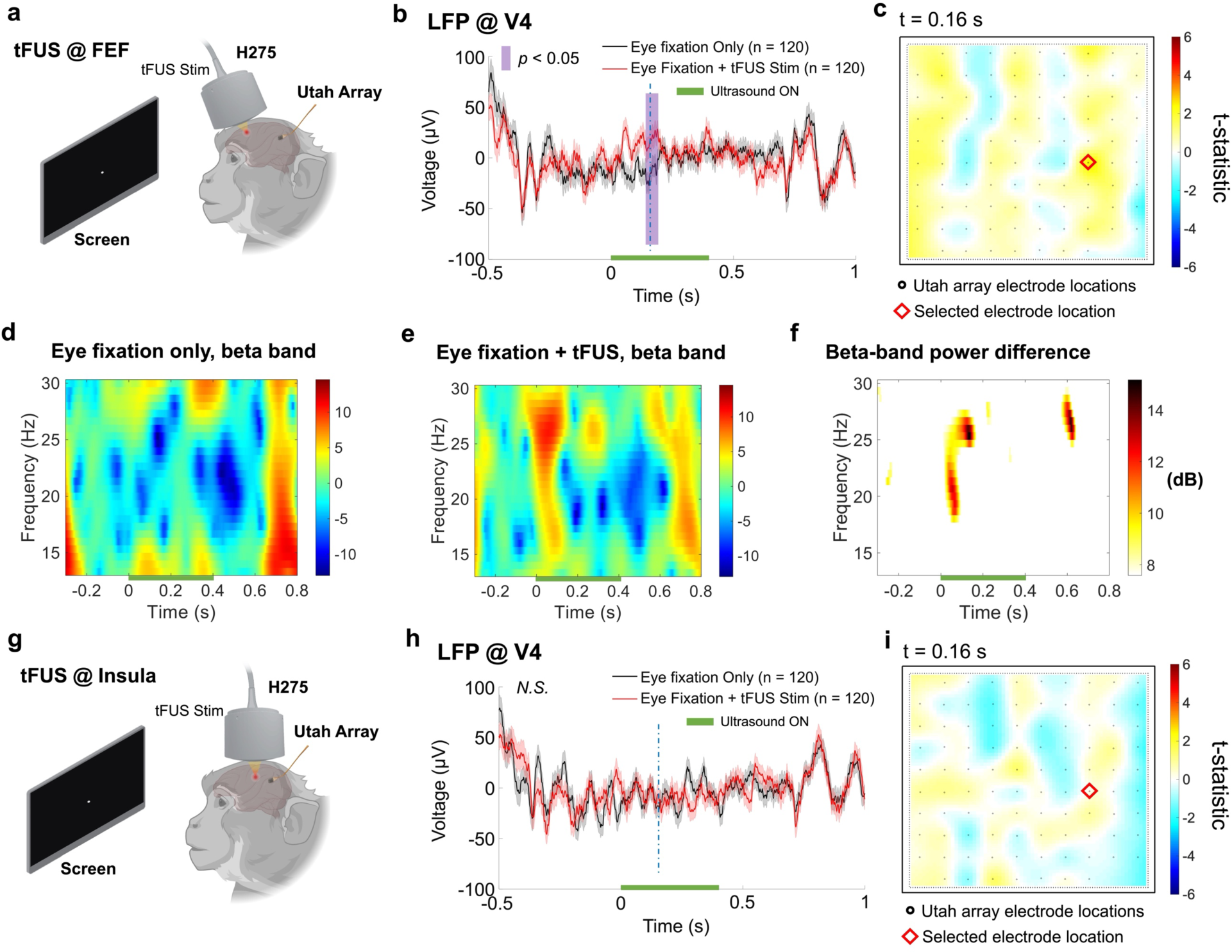
The time-frequency signature of tFUS-induced V4 activities without visual stimuli. **a**, The diagram of the experimental setup shows that a white dot appeared at the center of the screen and, after successful eye fixation was detected, tFUS stimulation was randomly (50% probability) delivered through H275 and targeted at FEF. No visual stimulus was presented on the screen. The LFPs at V4 were recorded through the Utah array. **b**. After repeating 120 trials with ultrasound stimulation, a single-channel LFP comparison showed a significantly different time segment versus the eye fixation-only condition. The vertical purple bar represents the significant difference (*p* < 0.05). The horizontal green bar shows the sonication period. **c**, A topographical map of t-statistics at 0.16 s (vertical blue dashed line in panel (**b**)) across the 96-channel recording sites over the V4. The red diamond represents the selected electrode location for the LFP waveforms in (**b**). **d**-**f**, The LFP beta band (13-30 Hz) contents in the fixation only condition (**d**) and in the tFUS stimulation condition (**e**), with the beta-band power difference shown in (**f**). **g**, Time-locked activation at V4 was absent when tFUS was targeted at the insula without visual stimuli. Following successful fixation, tFUS focus was steered and the same ultrasound stimulation (i.e., PRF: 3 kHz, CPP: 140, sonication duration: 400 ms, duty cycle: 60%) was randomly (50% probability) delivered to the insula. **h**, The LFP waveforms were derived from the fixation-only condition and the condition with tFUS stimulation (trial number: 120) at the same electrode site as the one in (**b**). Solid red/black lines indicate the mean, and pink/gray shaded areas indicate the S.E.M. *N.S.*, not significant. **i**, Another topographical map of t-statistics at 0.16 s (vertical blue dashed line in panel (**h**)) across the 96-channel recording sites over the V4 when only the tFUS stimulation was delivered to the insula. The red diamond represents the same selected electrode location as where the LFP waveforms in (**h**) were recorded. Statistics by cluster-based permutation test using Monte-Carlo estimates of the significance probabilities.

tFUS stimulation has been reported to induce auditory side effects via skull bone conduction^20^. In addition to our decoupled sham stimulation to rule out the airborne auditory effects, to exclude that the observed V4 activations were originated from the auditory pathway, we selected the insular cortex for tFUS stimulation. Although this is a deep cortical structure like FEF, it does not have any particularly strong connections to V4 and was, therefore, a suitable alternative stimulation location as a control. To target this deep cortical structure, we slightly adjusted the location of H275 towards the posterior and steered the focus to reach deeper (Δz = 9.85 mm). The task was the same as described above, with no visual stimulus (only a fixation dot at the center of the screen, Fig. 5g). When tFUS was directed onto the insular cortex, the steered transcranial ultrasound focus was measured at a lateral focal size of 2.48 mm and an axial focal size of 8.45 mm (-6 dB contour, Supplementary Fig. S1). To compensate for the increased steering depth, we slightly raised the transmission voltage of the Vantage system, thus increasing the transcranial spatial peak pressure to 1.14 MPa (Supplementary Fig. S1b). Even though we applied the same ultrasound temporal parameters as those administered to FEF (Fig. 5b-c), i.e., PRF: 3 kHz, duty cycle: 60%, LFP comparisons at the same marked electrode as in Fig. 5b-c in the time course confirmed as not significant (Fig. 5h-i). The vertical dashed lines in Fig. 5b and Fig. 5h indicate the 0.16 s time point.

## Discussion

In our study, we showed that ultrasound stimulation of frontal cortex can modulate the activity of neurons in visual cortex. Furthermore, we demonstrated that spatial steering of the ultrasound signal altered the effects on V4, consistent with the known anatomy of FEF and previous experiments with electrical stimulation. This observation was possible due to the increased spatial specificity and electronic beam steering achieved by the customized 128-element random array ultrasound transducer, permitting us to scan through the depth of FEF and further test the subregional specific neuromodulation effects on the visual-evoked potentials at V4 induced by tFUS at FEF.

Prior research shows that changing the ultrasound pulse repetition frequency can preferentially stimulate the excitatory or inhibitory neuronal populations in the somatosensory cortex^21^. In this previous work, the intracranial electrophysiological recordings were colocalized at the same brain region as the ultrasound stimulation target, and the recorded neuronal activities were characterized by the spiking rates during the sonication period. In the present work, we observed both enhancement and suppression of activity in LFPs and spiking in a site remote from the ultrasound stimulation. We found that the neuromodulation roles of tFUS in exciting or inhibiting the time course of LFP and single neuronal spiking activities during and/or after the sonication period depended on the applied PRF levels while using a constant burst duty cycle (Fig. 2d-g and Fig. 4b-d). Through this remote excitation and inhibition in the specific brain network of an awake, behaving animal model, we validated PRF’s importance in achieving different neuromodulation effects induced by tFUS.

Besides the critical role of the PRF, the higher burst duty cycle of tFUS also contributes to increased neural activations (Fig. 4d-i). Previously in an awake sheep model, the duty cycle of tFUS stimulation at motor cortex and thalamus was deemed a key factor of the excitatory or suppressive effect of tFUS on muscle activity^22^. Although our study targets a different neural pathway, the direct comparisons of tFUS parameter sets implemented in our work (i.e., changed PRFs with a constant duty cycle, and changed duty cycles with a constant PRF) further emphasize the parameter-dependent nature of tFUS neuromodulation.

With the improved spatial specificity of the ultrasound array, we set out to study the subregion-specific neuromodulation effects of tFUS by scanning through a specific cortical area, i.e., the frontal eye field, which is part of the cortical network controlling visual attention and eye movements. By gradually steering the ultrasound focus more ventrally into the sulcus of the FEF, more pronounced modulations to the visual cortical activities were seen (Fig. 3d-g). This location dependence of FEF modulation was observed previously with intracortical electrical stimulation^15^, and to our knowledge, this is the first evidence showing the subregional specific neuromodulation effects produced by noninvasive focused ultrasound stimulation.

By recording remotely from the site of ultrasound stimulation, we were able to eliminate numerous confounding factors that exist especially in physical electrode-based electrophysiological recordings (e.g., potential electrode vibration^23^). In this setting, delivering tFUS stimulation remotely was a rigorous way to study tFUS-induced effects in a well-defined brain network without being confounded by potential mechanical vibrations of electrodes, through either the artifacts presented in the recordings or electrode-vibration-induced neural effects.

Folloni et al. delivered transcranial ultrasound stimulation (PRF: 10 Hz) to the amygdala and anterior cingulate cortex in macaques, and their fMRI results revealed that the ultrasound stimulation caused the stimulated brain regions to decouple from their normally interconnected brain areas, demonstrating “a relatively focal and circumscribed impact” of ultrasound stimulation on the neural activity^6^. However, in our study, the remote inhibition and excitation of V4 demonstrate that this FEF-V4 connection is maintained and can be modulated with excitatory or inhibitory effects depending on a specific ultrasound parameter, i.e., PRF, administered onto FEF.

Therefore, our finding extends our understanding of ultrasound parameter control as a way to impact a specific neural pathway, and thus overall brain networks. Additionally, transcranial ultrasound stimulation at a PRF of 10 Hz^4,6^ warrants further investigation as a potential means to disrupt or alter FEF neuronal activity and thereby impact V4.

To validate whether the achieved remote neuromodulation by tFUS is specific to the brain target, we steered the ultrasound focus to a deeper cortical area, i.e., the insula, which does not have substantial connections to V4. Without presenting the visual stimuli on the screen, the tFUS stimulation at the insula (Fig. 5g) did not lead to any significant modulatory effects in the LFP waveforms (Fig. 5h) at V4. This control study confirmed that the tFUS neuromodulation was highly region specific, and the remote neural effects relied on the original stimulation site. Moreover, through this control study, we can also rule out the possibility that potential auditory side effects^24,25^, mainly induced by the tFUS^20^, could modulate the remote V4 neural activities.

Our study demonstrated that tFUS can produce nuanced neuromodulatory effects in cortical regions distant from the stimulation. Such stimulations on the animal subject were also observed as safe (see Supplementary Note 2: tFUS Safety Monitoring). In future work, it will be crucial to delve deeper into the nuanced behavioral alterations of such stimulation, including the trajectory and velocity of eye movements and the subtleties of visual perception, that result from administering tFUS with different parametric profiles and subregional targets. By analyzing behavior along with neural recordings, we can gain a clearer insight into the distant neuromodulatory effects of tFUS, and, in turn, advance its use as a powerful neuromodulatory technology.

## Data availability

The data are presented in the paper. Additional data will be made public through a data repository upon paper acceptance.

## Acknowledgments

This work was supported in part by NIH grants RF1NS131069, U18EB029354, and R01NS124564.

## Author Contributions

**Conceptualization**: K.Y., B.H., and M.S. **Methodology**: K.Y., M.S., S.S., and Y.N. **Data collection**: S.S., K.Y., Y.N., and E.C. **Formal analysis**: K.Y. **Investigation**: K.Y., M.S., S.S., and B.H. **Writing** – **Original draft**: K.Y. **Writing** – **Reviewing and editing**: K.Y., M.S., B.H., S.S., and E.C. **Supervision**: B.H., M.S.

