## Supplementary Information for "Transcranial Focused Ultrasound Remotely Modulates Extrastriate Visual Cortex with Subregion Specificity"

# of

### Supplementary Note 1: Structural Imaging

We acquired structural magnetic resonance imaging (MRI) and computed tomography (CT) images to build a subject-specific head model. Next, we segmented the head MRI and CT data to build specific brain and skull models, respectively. The mechanical properties of the brain and skull model were further assigned by referring to the database of IT'IS 4.0 (*The Foundation for Research on Information Technologies in Society*). The mass density, speed of sound, attenuation coefficient and characteristic acoustic impedance of the brain and skull tissues (cortical bone) are listed in Supplementary Table 1.

**Supplementary Table 1. Mechanical properties of the head model in computer simulations**

| Properties (Units) | Brain | Cortical Bone |
|---|---|---|
| Mass density (kg/m$^3$) | 1045.50 | 1908.00 |
| Speed of Sound (m/s) | 1546.32 | 3514.86 |
| Attenuation Coefficient (Np/m) | 4.28 | 38.19 |
| Characteristic Acoustic Impedance (MRayl) | 1.62 | 6.71 |

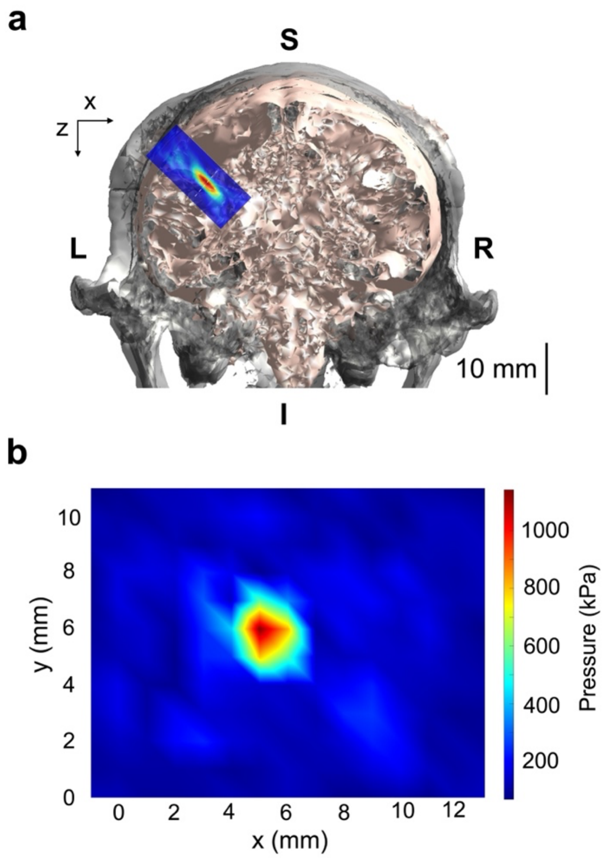


**Supplementary Figure S1.** *Ex-vivo* characterizations of transcranial focused ultrasound pressure field when targeting at insula. **a**, The axial profile of steered transcranial focused ultrasound beam measured using hydrophone scanning is superimposed onto the animal's coronal brain image, registering its location at the insula (RAS coordinates: -17.51, -9.9, 4.04). The ultrasound focus steering distance is calculated based on the RAS coordinates from the scalp. **b**, The lateral profile of the steered ultrasound focus targeting at the insula behind a skull sample. The presented lateral profile is the slice marked with white dashed line in panel (**a**). After penetrating through the skull bone, the spatial peak pressure was measured and estimated as 1.14 MPa from the *ex-vivo* scanning once the transcranial ultrasound focus is steered by the same distance calculated from the RAS coordinates.

**Supplementary Note 2: tFUS Safety Monitoring**

To assess tFUS safety, we measured the temperature using a circuit board (AD8495, Adafruit Industries, New York, NY, USA) connected to a thermocouple placed on the scalp at the sonication site during ultrasound stimulation. The observed temperature rise was less than 0.01°C. We also monitored the behaviors of the subject ten days before and ten days after the tFUS sessions. After repeating tFUS sessions over a week, the subject did not exhibit any abnormal gait, teeth grinding, repetitive motor activity, reluctance to stand, or lack of appetite that might have indicated an adverse impact. We also tracked average water earned in the task paradigm before and after tFUS and found the levels were comparable, indicating a stable behavioral state.
